# Stress fiber strain recognition by the LIM protein testin is cryptic and mediated by RhoA

**DOI:** 10.1101/2021.01.21.427693

**Authors:** Stefano Sala, Patrick W. Oakes

## Abstract

The actin cytoskeleton is a key regulator of mechanical processes in cells. The family of LIM domain proteins have recently emerged as important mechanoresponsive cytoskeletal elements capable of sensing strain in the actin cytoskeleton. The mechanisms regulating this mechanosensitive behavior, however, remain poorly understood. Here we show that the LIM domain protein testin is peculiar in that despite the full-length protein primarily appearing diffuse in the cytoplasm, the C-terminal LIM domains alone recognize focal adhesions and strained actin while the N-terminal domains alone recognize stress fibers. Phosphorylation mutations in the dimerization regions of testin, however, reveal its mechanosensitivity and cause it to relocate to focal adhesions and sites of strain in the actin cytoskeleton. Finally, we demonstrate activated RhoA causes testin to adorn stress fibers and become mechanosensitive. Together, our data show that testin’s mechanoresponse is regulated in cells and provide new insights into LIM domain protein recognition of the actin cytoskeleton mechanical state.

## Introduction

Cells depend on complex networks of both biochemical and mechanical signals which contribute to normal cellular physiology, tissue development and homeostasis (Lecuit et al., 2011; Iskratsch et al., 2014). While the general mechanisms of biochemical signaling are well defined, the mechanisms of mechanical signaling are less clear. Cells must recognize a variety of mechanical cues (e.g. tension, shape changes, pressure, stiffness), a behavior often referred to as mechanosensing, and convert those mechanical signals into biochemical signals, a process collectively called mechanotransduction (Paluch et al., 2015). These processes can influence a diverse array of critical functions including cell spreading (Oakes et al., 2018), force generation (Prager-Khoutorsky et al., 2011), proliferation (Rauskolb et al., 2014), migration (Das et al., 2015) and differentiation (Engler et al., 2006). Correspondingly, defects in mechanical signaling are associated with various diseases including muscular dystrophies and cardiomyopathies (Heydemann and McNally, 2007), osteoporosis (Hemmatian et al., 2017), asthma (Fabry and Fredberg, 2007), cancer progression and metastasis (Paszek et al., 2005).

The actin cytoskeleton and its associated binding proteins are key regulators of mechanical processes within and between cells (Ohashi et al., 2017; Blanchoin et al., 2014). Of the many diverse and dynamic architectures these proteins can form, two in particular play centralized roles in force transmission and mechanotransduction: focal adhesions (FA) and stress fibers (SF). FAs are complexes of ∼150 proteins which couple the actin cytoskeleton to the extracellular environment, transmitting forces between the cell and the extracellular matrix (Kanchanawong et al., 2010; Zaidel-Bar et al., 2007). Those forces are generated primarily by non-muscle myosin II motors which pull on the actin filaments in SFs, propagating the force to the FAs where they terminate (Schwarz and Gardel, 2012; Oakes and Gardel, 2014). SFs themselves are composed of ∼10-30 cross-linked actin filaments and can span tens of microns across the entire cell, providing an overall mechanical coherence to the cell (Tojkander et al., 2012; Cai and Sheetz, 2009; Svitkina, 2018).

As the combination of SFs and FAs serves as the primary conduit for force transmission in adherent cells, it is unsurprising that many proteins previously identified as mechanosensitive, such as alpha-actinin (Le et al., 2017), talin (Haining et al., 2016), vinculin (Goldmann, 2016), and integrins (Sun et al., 2016), are found in these structures. A common feature of these proteins is altered behavior under load, such as exposure of cryptic binding sites (Brown and Discher, 2009) or modified binding kinetics (Kong et al., 2009). An alternative and distinct mechanosensing mechanism, however, has been observed for the FA protein zyxin, which is induced to relocate to SFs under strain, whether through naturally occurring tears (Smith et al., 2011), optogenetically induced strain (Oakes et al., 2017) or when cells are placed under an externally applied cyclic stress (Hoffman et al., 2012; Yoshigi et al., 2005).

The family of proteins which are characterized by their LIM domains, consisting of two zinc-fingers, act as important scaffolds for protein-protein interactions (Smith et al., 2014; Sala and Ampe, 2018; Koch et al., 2012; Gill, 1995). Over the last two decades, many LIM domain-containing proteins (LDP) have, in addition to zyxin, been identified as mechanoresponsive cytoskeletal elements including paxillin (Smith et al., 2013), Hic-5 and CRP2 (Suzuki et al., 2005). More recently, it has been shown that several LDPs, including members of the zyxin, paxillin, Enigma and FHL families, share an evolutionarily conserved mechanism by which their LIM domains recognize mechanically strained F-actin (Sun et al., 2020; Winkelman et al., 2020). In all of these cases, their tandem LIM domains were necessary and sufficient for this mechanosensitive relocation. Remarkably, however, not all LDPs display mechanosensitive behavior and the mechanoresponse is not identical amongst LDPs. For instance, both zyxin and paxillin recognize SF strain sites (SFSS) but only zyxin relocates to SFs in response to cyclical stretch (Smith et al., 2013; Suzuki et al., 2005). This implies that LIM domain protein mechanosensitivity is specific and regulated. Dissecting these LDP-specific mechanosensitivity mechanisms and how they are regulated is necessary to better understand the molecular mechanisms underlying cellular mechanoresponse.

To probe this topic further, we investigated the mechanosensitivity of the multimodular LDP testin. Testin is expressed in the majority of tissues and consists of an N-terminal CR (Cysteine-rich), central PET (Prickle, Espinas, Testin) and three C-terminal LIM (Lin-11, Isl-1, Mec-3) domains (Boëda et al., 2011; Sala et al., 2017a; Tobias et al., 2001). While the exact function of testin remains tricky to identify, reduced expression has been observed in various tumors and cancer cell lines which, in conjunction with a number of animal studies, suggests a tumor suppressor function (Sarti et al., 2005; Drusco et al., 2005; Zhu et al., 2012; Gu et al., 2014). While testin has primarily been associated with the actin cytoskeleton, a proteomics-based interactome study additionally identified over 100 putative testin binding partners in a domain-specific manner, linking testin to adhesions, microtubules, endocytosis and nuclear receptor-mediated transcription among others, illustrating testin’s likely multifunctional nature (Sala et al., 2017b). Truncation variants consisting of either the N-terminal CR and PET domains or C-terminal LIM domains, have also been shown to localize to SFs and FAs, respectively (Garvalov et al., 2003; Sala et al., 2017b). In contrast, full-length testin is mainly distributed in the cytoplasm, consistent with its multi-conformational nature (Garvalov et al., 2003; Sala et al., 2017a). Together these findings suggest that the regulation of testin’s activity, and thus its mechanosensitivity, is likely mediated by its conformational state. We therefore hypothesized that testin is mechanosensitive under certain conditions and exploited its modular nature to investigate its potential role in recognizing local strain in the actin cytoskeleton.

## Results

### The LIM domains of testin, but not the full-length protein, recognize stress fiber strain sites

A subset of LIM domain proteins, including members of the FHL, zyxin and paxillin families, are known to recognize SFSSs via their LIM domains (Smith et al., 2011; Sun et al., 2020; Winkelman et al., 2020). To investigate whether the LIM domains of testin (LIM 1-2-3, Fig. 1A) display a similar mechanosensitive behavior, we engineered a series of truncations fused to GFP and co-expressed them with mApple-actin in human foreskin fibroblasts (HFF). LIM 1-2-3 localizes to FAs as previously reported (Fig. 1A, (Garvalov et al., 2003)). We then used a laser photoablation system to locally induce a SFSS, marked by a recoil and thinning of the SF (Fig. 1B, (Smith et al., 2011)). Potential recruitment was monitored by tracking the GFP signal over time using a mask based on the brightest pixels in the region surrounding the ablation that increased the most in intensity compared to their pre-ablation state (Figs. 1B,C, S1). The same pixels were then used to monitor the intensity of actin in those regions. Following photoablation, we observed a rapid increase in the GFP signal within seconds as LIM 1-2-3 relocated to the induced SFSS and a concomitant loss of actin intensity as the SF was damaged (Figs. 1B,C, Movie S1). Following LIM 1-2-3 recruitment, the actin signal subsequently recovered within minutes, indicating SF repair (Smith et al., 2011), before it finally plateaued. To verify that laser-induced damage actually results in the induction of a SFSS, we combined our laser photoablation approach with traction force microscopy and show that the ablated SFs remain under tension while LIM 1-2-3 is recruited (Fig. S2, Movie S2).

**Figure 1.**
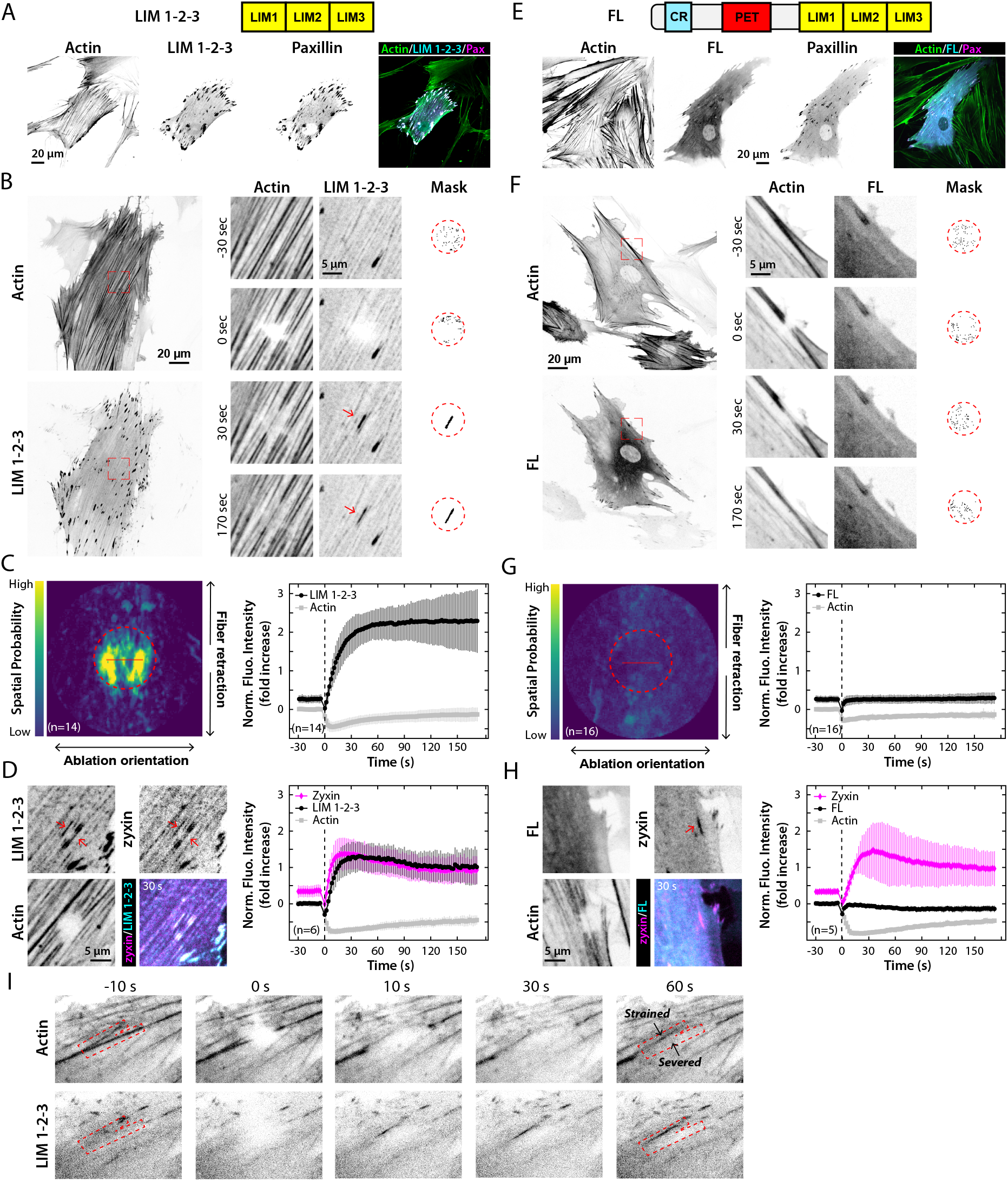
The LIM domains of testin, but not full-length, recognize stress fiber strain sites. **A**,**E)** Domain organization and subcellular localization of the LIM domains of testin (LIM 1-2-3) and full-length testin (FL) in HFFs. GFP-LIM 1-2-3 localizes strongly to FAs (A) whereas GFP-FL displays a diffuse cytoplasmic and weak FA localization (E). GFP-testin, mApple-paxillin and SiR-Actin channels are shown. **B**,**F)** HFFs co-expressing mApple-actin and either GFP-LIM 1-2-3 (B) or GFP-FL testin (F). Magnified inset images (red boxed regions) display the localization of each construct during the timelapse. Red arrows indicate relocation of LIM 1-2-3 to SFSSs (Movie S1). Masks show the pixels in the region surrounding the ablation (red dashed circle) that increased the most in intensity compared to their pre-ablation state. **C**,**G)** Spatial probability of testin recruitment (left) and average fluorescence intensity traces with standard deviation of the testin and actin signals (right) in the region of strain. The brightest LIM 1-2-3 signal is located in the central region of strain (C, dashed circle) whereas the brightest FL signal is distributed randomly (G). Red line represents the 5 µm laser line used to photo-induce SFSSs. Following SFSS induction, a rapid and large increase of the LIM 1-2-3 signal (C) but no change of the FL testin signal was detected in the region of strain (G). **D**,**H)** Experiments shown in C and G were repeated in the presence of the SFSS marker mCherry-zyxin, and SiR-Actin (Movie S3). Increased LIM 1-2-3 fluorescence intensity in the region of strain and colocalization with zyxin at SFSSs (D, red arrows). FL signal remained unchanged after ablation and did not colocalize with zyxin (H). I) HFFs expressing mApple-actin and GFP-LIM 1-2-3 during ablation of Y-shaped SFs are shown (red dashed box). LIM 1-2-3 relocates to the strained SF but not the severed SF (Movie S5).

To assess the spatial distribution of the LIM 1-2-3 localization, we aligned the masks to the angle of ablation, summed each experimental group over time, and averaged them to create a spatial probability map (Fig. S1D). This map clearly shows the brightest LIM 1-2-3 signal is located in the center of the mask stack where the ablation occurred (red line, Fig. 1C) and where the region of strain is located (dashed circle, Fig. 1C). To confirm that this is a SFSS, we assessed relocation of LIM 1-2-3 in the presence of mCherry-conjugated zyxin, a known SF strain sensor (Smith et al., 2011), and used SiR-Actin to visualize the SFs. Masks in these experiments were created based on regions where zyxin increased in intensity. Immediately following strain induction, both zyxin and LIM 1-2-3 relocated to and colocalized at SFs with similar kinetics, confirming the LIM domains of testin recognize actual SFSSs (Fig. 1D, Movie S3). Similar results were seen in zyxin-null MEFs (Fig. S3, Movie S4), indicating that relocation of LIM 1-2-3 to SFSSs is independent of zyxin.

In contrast to its LIM domains, full-length (FL) testin is mainly cytoplasmic with only a small subpopulation localizing to FAs (Fig. 1E). When a SFSS was induced, we did not observe recruitment of FL testin (Figs. 1F,G, Movie S1) and instead saw a rapid recovery of fluorescence intensity consistent with diffusion. Correspondingly, the spatial probability of recruitment map shows a random distribution and no regions of local enrichment in the central region of strain (Fig. 1G). When this experiment was repeated using zyxin as a control, we observed a rapid relocation of zyxin, but not testin, to sites of strain. This confirms that in our steady-state conditions FL testin is unable to recognize SFSSs (Fig. 1H, Movie S3).

### The LIM domains of testin recognize strained, but not severed, SFs

Previous work has demonstrated the presence of free barbed ends in naturally occurring spontaneous SFSSs (Smith et al., 2011). This raises the possibility that the LIM domains of testin associate with these barbed ends in SFSSs in addition to recognizing strained actin fibers. To test this, we increased the local laser power to completely sever SF bundles and assessed whether testin’s LIM domains recognized free actin filament ends in the absence of strain. Fig. 1I shows two SFs arranged in a Y-shape (red box), where the bottom fiber is completely severed, confirmed by the complete loss of the actin signal, and the top fiber becomes strained. Recruitment of LIM 1-2-3 only occurred on the strained fiber, and not along the retracting fiber with the free ends (Fig. 1I, Movie S5). This demonstrates that testin’s LIM domains do not recognize barbed actin filament ends.

### A single LIM domain is sufficient to recognize SFSSs

To further dissect the mechanoresponsive regions within testin’s LIM domains, we investigated the ability of the individual LIM domains (LIM 1, LIM 2 and LIM 3; Fig. 2) to recognize SFSSs. Previous research has suggested that multiple LIM domains are required for mechanosensitivity (Sun et al., 2020; Winkelman et al., 2020). Surprisingly, not only did each individual LIM domain of testin display FA localization (Figs. 2A, 2D, 2G), they also relocated to SFSSs (Figs. 2B,C,E,F,H,I, J,K; Movie S6). In order to compare the recruitment magnitude and kinetics to SFSSs of testin’s LIM domains, we fit an exponential recovery curve to the average traces of the various constructs (Fig. S4) and calculated the t_1/2_, amplitude and plateau maximum of the exponential fit (Fig. 2L; Table S1). Based on these values, the recruitment kinetics of LIM 1-2-3 are very similar to those of the individual LIM domains. Of the individual LIM domains, LIM 2 showed the greatest recruitment, though this recruitment remained slightly less than LIM 1-2-3 (Table S1, Figs. 2K,L). These data illustrate that each of testin’s LIM domains recognize SFSSs and that their mechanosensitivity is blocked in the full-length protein.

**Figure 2.**
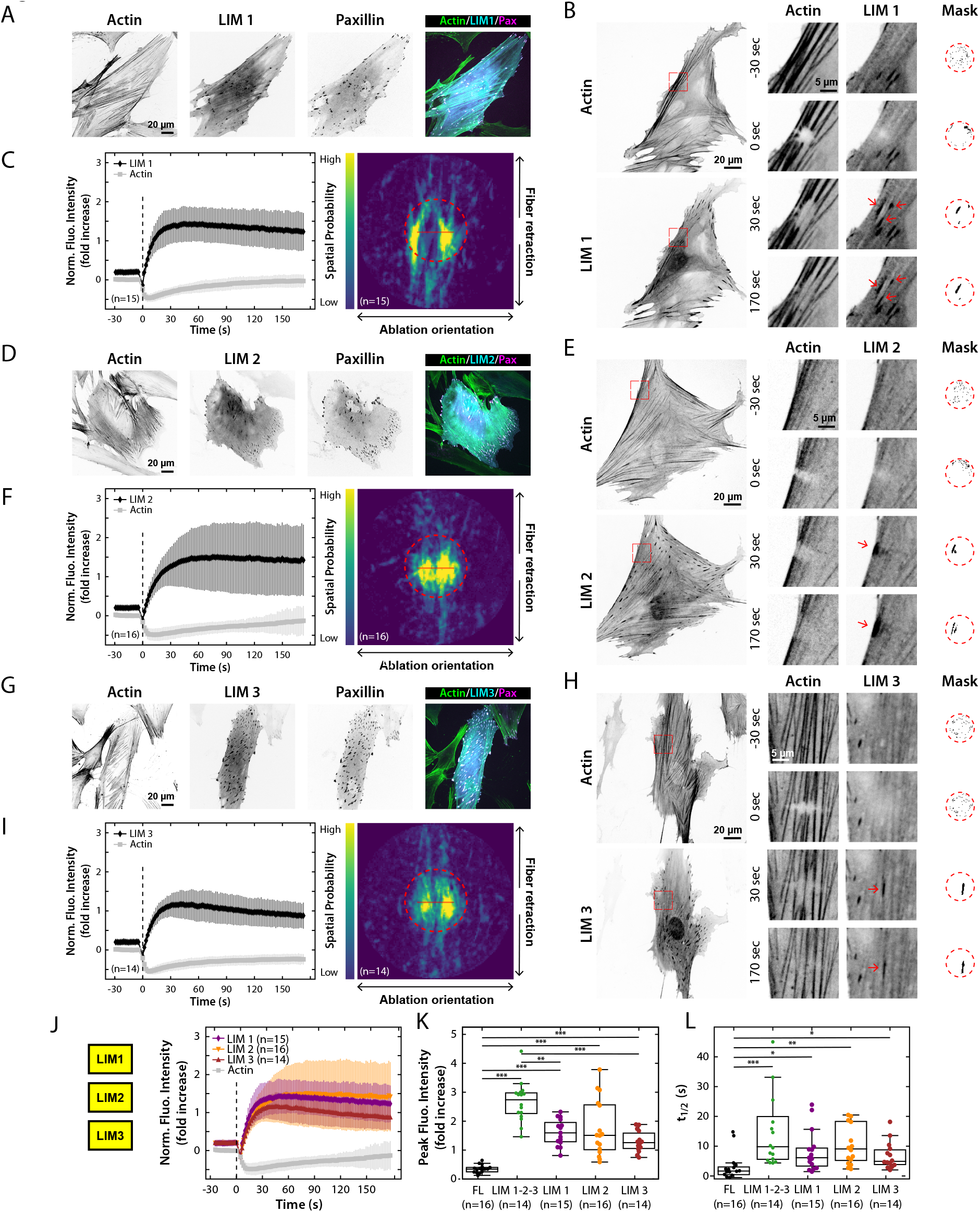
The individual LIM domains of testin are sufficient to recognize stress fiber strain sites. **A, D, G)** Subcellular localization of the individual LIM domains of testin in HFFs. All LIM domains localize to FAs. GFP-testin, mApple-paxillin and SiR-Actin channels are shown. **B, E, H)** HFFs expressing mApple-actin and GFP-LIM 1 (B) GFP-LIM 2 (E) or GFP-LIM 3 (H). Magnified inset images (red boxed regions) display the localization of each construct during the timelapse demonstrating relocation to SFSSs of each individual LIM domain (red arrows, Movie S6). Mask images show the pixels in the region surrounding the ablation (red dashed circle) that increased the most in intensity compared to their pre-ablation state. **C, F, I)** Average fluorescence intensity traces and standard deviation of the LIM and actin signals (left) and spatial probability of LIM recruitment (right) in the region of strain. Following SFSS induction, a rapid and large increase of all LIM signals was detected in the region of strain (left) and the brightest LIM signals were located in the central region of strain (right, dashed circles). Red line represents the 5 µm laser line used to photo-induce SFSSs. **J)** Average fluorescence intensity traces and standard deviation of testin’s individual GFP-coupled LIM domains and mApple-actin signals in the region of strain indicating recruitment to SFSSs of each LIM domain. **K)** Distribution of the peak fluorescence intensities after SFSS induction per testin variant. **L)** Recruitment T_1/2_ after SFSS induction per testin variant. *: p≤ 0.05., **: p≤ 0.01, ***: p≤ 0.001. Dashed vertical lines in C, F, I and J indicate the ablation timepoint.

### The N-terminal and C-terminal halves of testin recognize different mechanical states of F-actin

We next tested whether the N-terminal half of testin (NT), containing the CR and PET domains (Fig. 3A), is capable of recognizing SFSSs. Despite an initial strong SF localization (Fig. 3A), we did not observe any obvious enrichment of NT at SFSSs (Figs. 3B,C; Movie S1). However, quantification of the fluorescence intensity in the masked region of strain did reveal a slight increase of the NT signal (Figs. 3D,E). Upon closer examination we noticed that the actin signal simultaneously increased in these regions, opposite to what is expected for a SFSS (Fig. 3D). Examination of the spatial distribution of the recruitment revealed that the brightest NT signal was located outside of the central region of strain along the recoil axis of the actin (Fig. 3C) and was smaller in magnitude than the LIM 1-2-3 signal (Fig. 3E). Together these data indicate that, unlike LIM 1-2-3, the increase in fluorescence of NT arises from a condensation of severed actin filaments and not from additional recruitment (Figs. 3C,D). Therefore, in contrast to its LIM domains, the N-terminal half of testin is unable to bind SFSSs.

**Figure 3.**
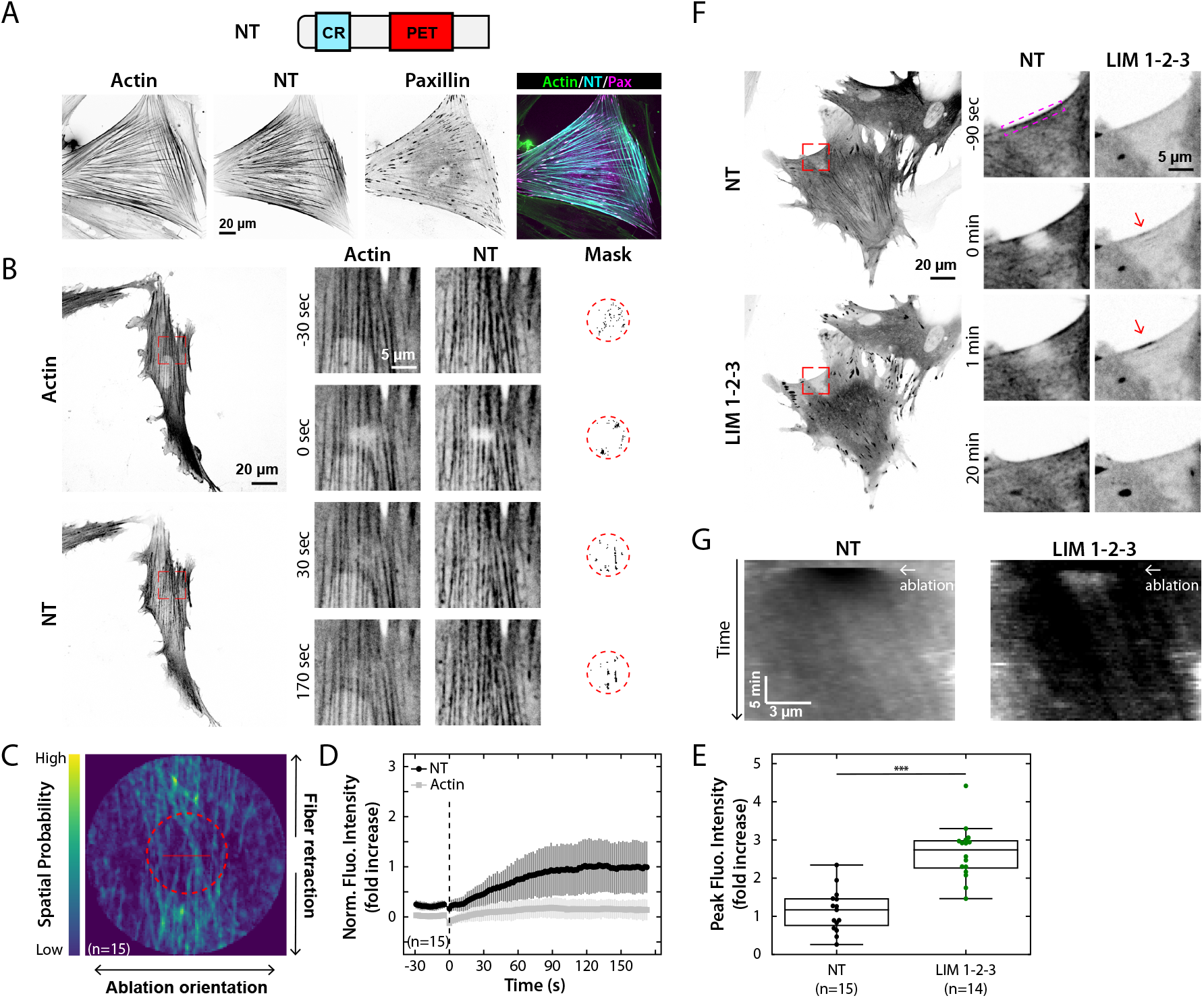
The N-terminal domains and C-terminal LIM domains of testin recognize different mechanical states of SFs. **A)** Domain composition and subcellular localization of the N-terminal domains of testin (NT). GFP-NT localizes strongly to SFs. GFP-NT, mApple-paxillin and SiR-Actin channels are shown. **B)** HFFs co-expressing mApple-actin and GFP-NT. Magnified inset images (red boxed regions) display the localization of NT during the timelapse (Movie S1). Masks show the pixels in the region surrounding the ablation (red dashed circle) that increased the most in intensity compared to their pre-ablation state. **C)** Spatial probability of NT recruitment in the region of strain shows NT is primarily recruited outside the central region of strain (red dashed circle). Red line represents the 5 µm laser line used to photo-induce SFSSs. **D)** Average fluorescence intensity trace and standard deviation of the NT and actin signals in the region of strain. Dashed vertical line indicates the ablation timepoint. **E)** Distribution of the peak fluorescence intensities after SFSS induction for NT and LIM 1-2-3. ***: p≤ 0.001. **F)** HFFs co-expressing GFP-LIM 1-2-3 and mApple-NT. Magnified inset images (red boxed regions) display the localization of each construct during timelapse showing recruitment of GFP-LIM 1-2-3 to SFSSs (red arrows) and absence of colocalization of GFP-LIM 1-2-3 and mApple-NT (Movie S7). **G)** Kymograph along the magenta boxed SF in F showing recruitment of LIM-1-2-3 to the SFSS following ablation and simultaneous dissociation of NT in the same region. Arrows indicate the ablation timepoint.

To further illustrate this, we co-expressed LIM 1-2-3 and NT and assessed whether the two halves of testin colocalized at all following induction of a SFSS (Fig. 3F, Movie S7). To ensure full repair of the ablated SF, we monitored their localization over a time period of 30 min (Figs. 3F,G). Prior to SFSS induction, LIM 1-2-3 localized to FAs whereas NT localized to SFs as mentioned above (Figs. 3F,G). After SFSS induction, LIM 1-2-3 was immediately recruited to the strain site (arrows, Figs. 3F,G) while NT simultaneously dissociated from the strain region (Figs. 3F,G). NT remained bound to SFs in the regions flanking the strain site. As the SFSS was repaired over time, LIM 1-2-3 dissociated from the ablated SF and NT reassociated with the repaired actin fibers (Figs. 3F,G). The absence of colocalization of NT and LIM 1-2-3 at either SFs or SFSSs demonstrates that the two halves of testin recognize different mechanical states of F-actin.

### Tyrosine mutations in the PET and LIM 1 domains drive testin to SFSSs and FAs

Given the multi-conformational nature of testin (Garvalov et al., 2003; Sala et al., 2017a), it is possible that the inability of FL testin to recognize SFSSs results from a conformational state that makes the LIM domains inaccessible. Previous work demonstrated that tyrosine 288 (Y288) in the first LIM domain is structurally important for testin dimerization since mutating this tyrosine to alanine strongly inhibits dimerization (Sala et al., 2017a). Therefore, we hypothesized that mutating Y288 to alanine would allow testin to recognize strain in the actin cytoskeleton by potentially adopting a more favorable conformation.

To investigate this, we co-expressed a GFP-Y288A mutant of testin (Y288A; Fig. 4A) along with actin in HFFs and assessed its ability to relocate to SFSSs (Movie S8). Induction of SFSSs resulted in a moderate but obvious recruitment of Y288A to the site of strain (Fig. 4A). The spatial probability map also revealed a preference for recruitment in the region of strain (Fig. 4B). To further investigate the potential regulatory role of Y288 in determining testin’s mechanosensitivity, we created a phosphomimetic glutamic acid (Y288E) and non-phosphorylatable phenylalanine (Y288F) mutants of testin and assessed their ability to recognize SFSSs (Figs. 4C,D,E,F; Movie S8). Similar to Y288A, Y288E was capable of recognizing SFSSs (Figs. 4C,D,G,H). Conversely, the Y288F mutant did not display this mechanosensitive behavior (Figs. 4E-H). These data illustrate that specific mutations of Y288 in the first LIM domain can promote testin’s mechanosensitivity.

**Figure 4.**
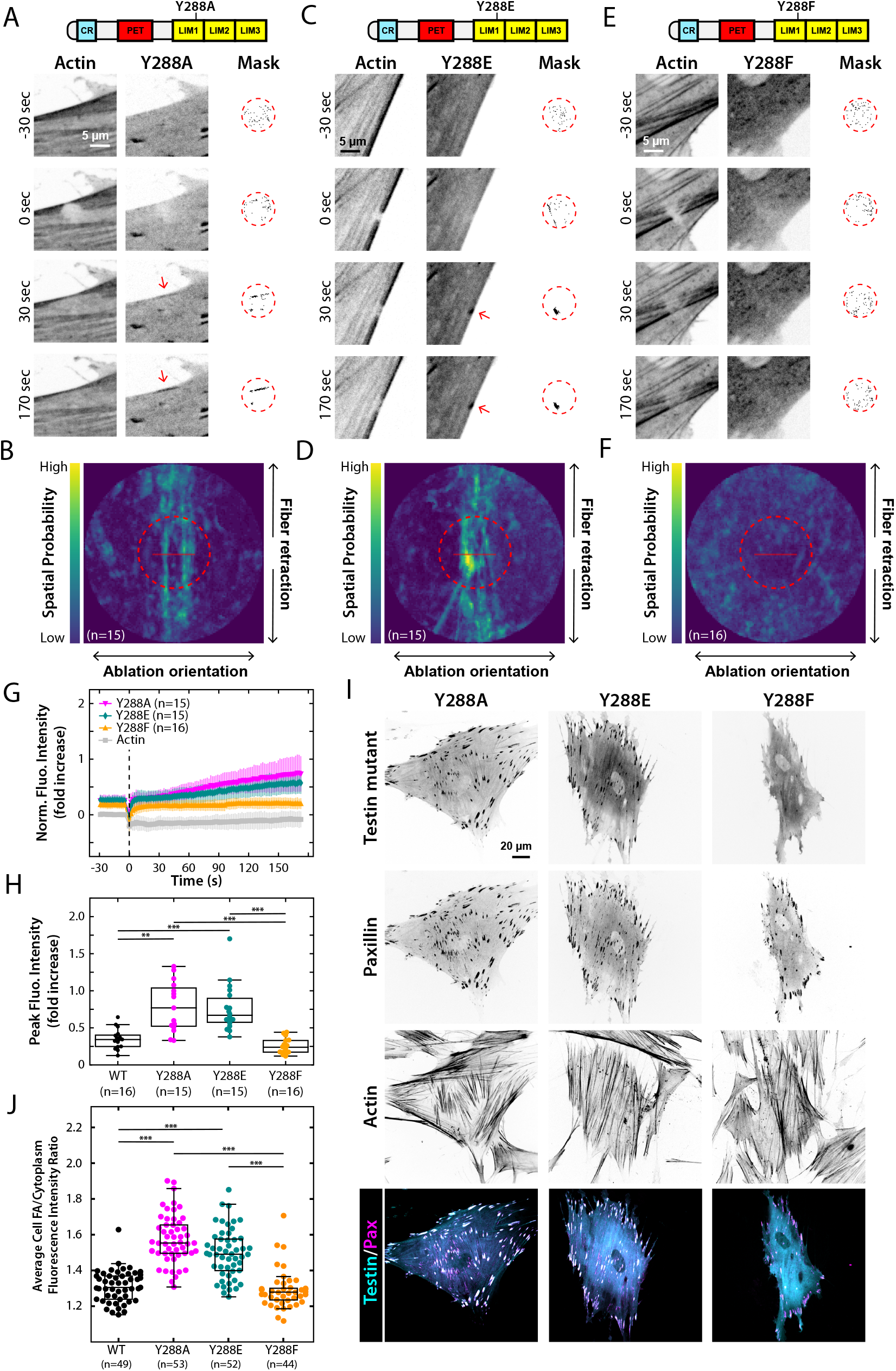
The Y288A and Y288E mutants of testin recognize SFSSs. **A, C, E)** HFFs co-expressing mApple-actin and either GFP-Y288A (A) GFP-Y288E (C) or GFP-Y288F (E). Magnified inset images (red boxed regions) display the localization of each construct during the timelapse demonstrating relocation to SFSSs of both the Y288A and Y288E mutants (A and C, red arrows), but not the Y288F mutant (E), (Movie S8). Masks show the pixels in the region surrounding the ablation (red dashed circle) that increased the most in intensity compared to their pre-ablation state. **B, D, F)** Spatial probability of recruitment of the Y288 testin mutants in the region of strain. The brightest Y288A and Y288E signals are located in the central region of strain (B and D, dashed circle) whereas the brightest Y288F signal is distributed randomly (F). Red lines represent the 5 µm laser line used to photo-induce SFSSs. **G)** Average fluorescence intensity traces and standard deviation of the Y288 testin mutants and actin signals in the region of strain indicating recruitment of the Y288A and Y288E mutants, but not the Y288F mutant, to SFSSs. Dashed vertical line indicates the ablation timepoint. **H)** Distribution of the peak fluorescence intensities after SFSS induction for WT testin and the Y288 mutants. **: p≤ 0.01, ***: p≤ 0.001. **I)** Subcellular localization in HFFs of the Y288 mutants. The mechanosensitive Y288A and Y288E mutants strongly localize to FAs whereas the non-mechanosensitive Y288F mutant displays a diffuse cytoplasmic localization and only weak FA localization. GFP-testin, mApple-paxillin and SiR-Actin channels are shown. **J)** Distribution of the average cellular FA/cytoplasmic fluorescence intensity ratio per testin variant. ***: p≤ 0.001.

The observation that Y288A/E mutations drive testin to SFSSs implies the accessibility of the LIM domains is enhanced in these mutants. Since the truncated LIM domains of testin, but not FL testin, strongly localize to FAs (Fig. 1A,E), we hypothesized that the mechanosensitive Y288A/E mutants would display increased FA localization. To investigate this, we co-expressed either Y288A or Y288E with paxillin, another FA protein, and assessed their FA localization (Fig. 4I). To quantify FA localization, we calculated the ratio of the fluorescence intensity in FAs and the cytoplasmic region surrounding the FAs for every FA in a given cell (Figs. 4J,S5). In agreement with the results obtained from the laser photoablation assay, we found that FA localization of the mechanosensitive Y288A/E mutants was significantly increased compared to the non-mechanosensitive Y288F mutant (Fig. 4J). The Y288F mutant displayed a FA localization that was comparable to the WT protein (Fig. 4J), which is consistent with their inability to recognize sites of SF strain.

Testin forms dimers through interactions involving its PET domain and its first two LIM domains (Sala et al., 2017a). Since specific mutations of Y288, located in the dimerization region of the protein, promote testin’s mechanosensitivity, we reasoned that other tyrosines in those dimerization regions might similarly affect its mechanosensitivity. We therefore investigated tyrosine 111 (Y111), located in the PET domain, by creating non-phosphorylatable alanine (Y111A) and phenylalanine (Y111F) mutants as well as a phosphomimetic glutamic acid (Y111E) mutant (Figs. 5A,B,C,D,E,F; Movie S8). Interestingly, we observed a similar behavior for the Y111 mutants with clear recruitment of Y111A (Figs. 5A,B,G-H) and Y111E (Figs. 5C,D,G-H) mutants to SFSSs but not the Y111F mutant (Figs. 5E-H). In agreement with the results from the Y288 mutants, FA localization of the mechanosensitive Y111A/E mutants was significantly increased compared to the non-mechanosensitive Y288F mutant (Fig. 5J). Our quantitative analysis further revealed that FA localization (Fig. 5J) of the Y111A and Y111E mutants as well as their relocation to sites of SF strain (Figs. 5G-H) was stronger than what we observed for the corresponding mechanosensitive Y288 mutants (Figs. 4G-H,J), suggesting these mutants recognize FAs and SFSSs with different affinities (Fig. S4;Table S1). Together, these results demonstrate that mutations in the dimerization regions of testin affect its ability to recognize strain in the actin cytoskeleton, and that the magnitude of the effect is dependent on the nature of the mutation.

**Figure 5.**
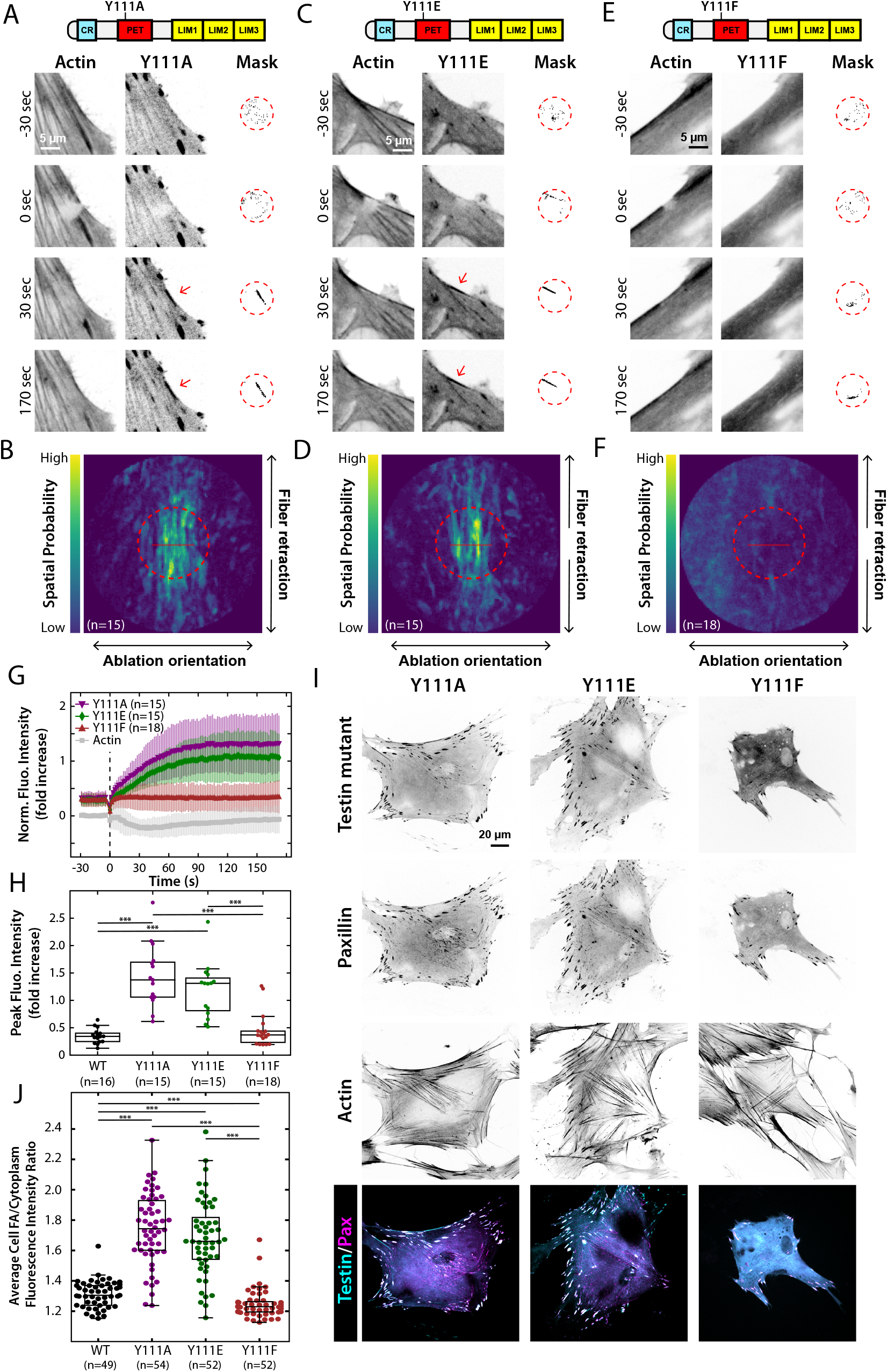
The Y111A and Y111E mutants of testin recognize SFSSs. **A, C, E)** HFFs co-expressing mApple-actin and either GFP-Y111A (A) GFP-Y111E (C) or GFP-Y111F (E). Magnified inset images (red boxed regions) display the localization of each construct during the timelapse demonstrating relocation to SFSSs of both the Y111A and Y111E mutants (A and C, red arrows), but not the Y111F mutant (E), (Movie S8). Masks show the pixels in the region surrounding the ablation (red dashed circle) that increased the most in intensity compared to their pre-ablation state. **B, D, F)** Spatial probability of recruitment of the Y111 testin mutants in the region of strain. The brightest Y111A and Y111E signals are located in the central region of strain (B and D, dashed circle) whereas the brightest Y111F signal is distributed randomly (F). Red lines represent the 5 µm laser line used to photo-induce SFSSs. **G)** Average fluorescence intensity traces and standard deviation of the Y111 testin mutants and actin signals in the region of strain indicating recruitment of the Y111A and Y111E mutants, but not the Y111F mutant, to SFSSs. Dashed vertical line indicates the ablation timepoint. **H)** Distribution of the peak fluorescence intensities after SFSS induction for WT testin and the Y111 mutants. ***: p≤ 0.001. **I)** Subcellular localization in HFFs of the Y111 mutants. The mechanosensitive Y111A and Y111E mutants strongly localize to FAs whereas the non-mechanosensitive Y111F mutant displays a diffuse cytoplasmic localization and only weak FA localization. GFP-testin, mApple-paxillin and SiR-Actin channels are shown. **J)** Distribution of the average cellular FA/cytoplasmic fluorescence intensity ratio per testin variant. ***: p≤ 0.001.

### Activated RhoA promotes SF localization of testin

The behavior of the various truncations and tyrosine mutants in comparison to WT FL testin, implies that its mechanosensitivity must be regulated. Since local strain induction at SFs is not sufficient for testin to recognize these sites, we hypothesized that perturbing upstream signaling regulating the actin cytoskeleton at the cellular scale, might promote testin’s mechanosensitivity. Since RhoA is known to modulate the overall cellular contractility and actin cytoskeleton organization (Ridley and Hall, 1992; Oakes et al., 2017), we transfected cells with mApple-conjugated FL testin in conjunction with either a constitutively active RhoA (CA-RhoA) or dominant negative RhoA (DN-RhoA) variant and assessed the localization of testin under different levels of cytoskeletal tension (Fig. 6A). Cells expressing the CA-RhoA variant were more contractile (Fig. 6B) and 90% of these cells exhibited SF localization of testin (Figs. 6A,C), compared to 26% of the control cells where testin was primarily cytoplasmic (Figs. 6A,C). Correspondingly, in only 17% of the DN-RhoA expressing cells, testin localized to SFs and was mainly distributed in the cytoplasm (Figs. 6A,C).

**Figure 6.**
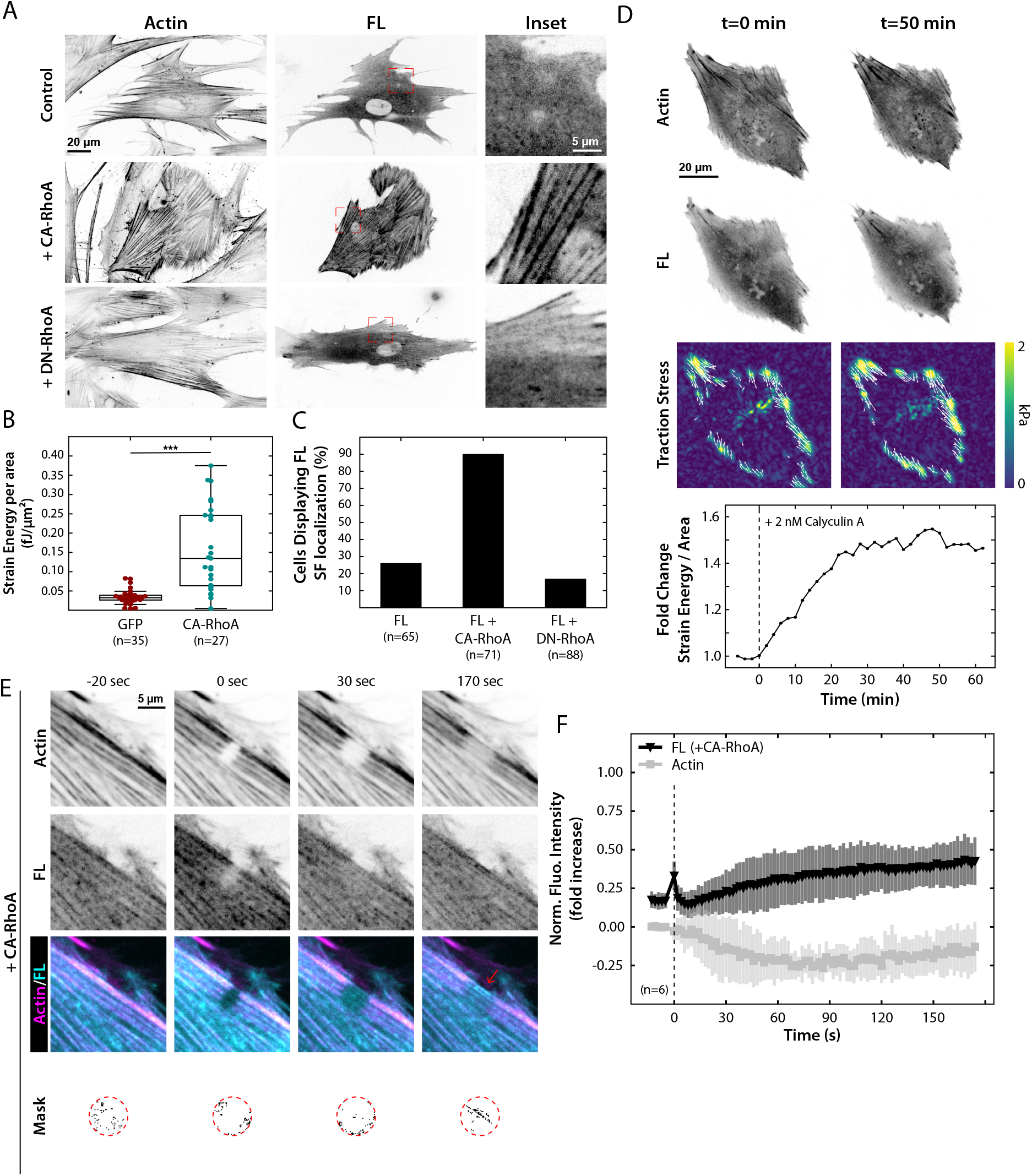
RhoA promotes SF localization of testin. **A)** HFFs expressing mApple-FL testin and either CA GFP-RhoA or DN GFP-RhoA. In the presence of CA GFP-RhoA, mApple-FL testin relocates to SFs (magnified insets). **B)** Measurement of strain energy per area is shown for both the control (GFP) and CA-RhoA expressing cells. In the presence of CA-RhoA, the overall contractility of the cell is increased. ***: p≤ 0.001. **C)** Percentage of HFFs displaying SF localization of mApple-FL testin in the absence or presence of either CA-RhoA or DN-RhoA. **D)** Representative images of HFFs expressing GFP-actin and mApple-FL testin and their corresponding traction stresses before and after addition of 2nM Calyculin A. Traction stress vectors (white arrows) are only shown for regions greater than 500 Pa. Calyculin A increases cellular contractility (vertical dashed line) as indicated by the increase in traction stresses and relative change in strain energy per area. **E)** HFFs co-expressing mApple-FL testin and CA GFP-RhoA during an ablation. SFs were stained with SiR-Actin. Following strain induction, FL testin relocated to SFSSs (red arrow, Movie S9). **F)** Average fluorescence intensity traces and standard deviation of FL testin and actin in the region of strain indicating recruitment of FL testin to SFSSs. Dashed vertical line indicates the ablation timepoint.

To investigate whether increased SF localization of testin is driven by RhoA or merely the effect of increased cellular contractility and tension, we monitored testin localization in cells treated with the drug calyculin A to increase cellular contractility without stimulating RhoA activity (Chartier et al., 1991; Lemmon et al., 2009; Stricker et al., 2011). Traction force microscopy analysis demonstrates the cells become more contractile after calyculin A treatment (Fig 6D). Testin localization, however, remained unaffected (n=14 cells examined) illustrating that increased cellular contractility is not sufficient to drive the protein to SFs and that upstream RhoA signaling is required for testin relocation to SFs (Fig 6A-D).

Given that activated RhoA affects testin localization, we next examined whether elevated RhoA activity also enables testin to recognize SF strain. To test this, we repeated our laser photoablation experiment with FL testin in cells expressing CA-RhoA (Figs. 6E,F, Movie S9). In contrast to the control cells (Figs. 1F-G), we saw recruitment of FL testin to the site of strain (Fig. 8E) and an increase in fluorescence intensity of testin over time that corresponded with a simultaneous decrease in the actin intensity in those regions (Fig. 6F). Combined, these results indicate that elevated RhoA activity promotes testin’s SF localization and mechanosensitivity.

## Discussion

Our data reveal that the mechanosensitivity of the LDP testin is highly regulated, likely through modification of its biochemical and/or conformational state, allowing cells to potentially control its spatiotemporal behavior through signaling molecules such as RhoA. Over the last decade, it has become more apparent that the mechanical state of filamentous actin (tension, curvature, torsion) affects the binding and activity of many actin-binding proteins (Jégou and Romet-Lemonne, 2021; Zimmermann et al., 2017). Recent work (Sun et al., 2020; Winkelman et al., 2020) has suggested that a conserved mechanism in LIM domains confers mechanosensitivity to proteins from the FHL, zyxin and paxillin families among others to sense strained actin filaments. The LDP testin appears unique in that the FL protein resides primarily in the cytoplasm, while its N-terminal half recognizes SFs and its C-terminal half recognizes SFSSs and FAs (Figs. 1-3). Although the functional consequences of this unique feature are currently unknown, we speculate that testin acts as a bifunctional scaffold protein in mechanically stressed actin-rich structures.

Recent works comparing LDPs have shown that there exists a large diversity in their LIM domain sequences and have suggested that multiple successive LIM domains are required to recognize SFSSs (Sun et al., 2020; Winkelman et al., 2020). We show, however, that each of the individual LIM domains of testin can recognize SFSSs on their own in cells (Fig. 2). This discrepancy between testin and other LDPs is likely due to differences in the sequence of testin’s LIM domains. Sun et al. have proposed that a conserved phenylalanine in LIM domains is vital for their mechanosensitivity (Sun et al., 2020), but only one of testin’s LIM domains (LIM 3) contains this amino acid in the conserved place (Sala et al., 2017a). This seeming contradiction could have a number of potential explanations. First, and most excitingly, different LDPs could use different mechanisms to recognize SFSSs. Alternatively, the relative amount of strain in the actin filaments could differentially affect their conformation and hence the ability of a LIM domain to associate with them. In other words, the strain we induce in actin filaments by using a laser might be slightly different compared to the strain induced through natural tears and ruptures. Lastly, we cannot rule out that testin’s individual LIM domains are indirectly associating with strained actin via recruitment by other SFSS proteins. The LIM domains of testin have been shown to interact with zyxin, VASP and alpha-actinin (Garvalov et al., 2003; Sala et al., 2017b), which are all found in SFSSs (Smith et al., 2011). This last scenario, however, seems unlikely since we find that LIM 1-2-3 recruitment to SFSSs is independent of zyxin (Fig. S3) and, more importantly, that the recruitment kinetics of the individual LIM domains of testin are all comparable to the recruitment kinetics of LIM 1-2-3 (Fig. S4). Ultimately, *in vitro* and structural studies of individual LIM domains bound to strained actin will likely be necessary to elucidate the exact mechanisms. Finally, the differences in magnitude of SFSS recognition of the individual LIM domains of testin and the LIM 1-2-3 variant are likely a result of an additive contribution as suggested previously for zyxin’s LIM domains (Winkelman et al., 2020).

While the N-terminal half of testin clearly localizes to SFs, our data illustrate that it is not mechanosensitive on its own (Fig. 3). We hypothesize that it might play a role in maintaining SF homeostasis in conjunction with the LIM domains. Certain actin-binding proteins have indeed been shown to form co-complexes with either the LIM domains (e.g. Ena/VASP proteins, (Garvalov et al., 2003; Boëda et al., 2007; Sala et al., 2017b)) or N-terminal domains (e.g. calponin-2, (Sala et al., 2017b)) of testin, showing that both halves could contribute to the recruitment of these proteins. Future studies will be required to investigate the potential function of testin in maintaining the mechanical integrity of SFs.

The numerous point mutations which confer mechanosensitivity to testin suggest that this function of the protein can be regulated. High throughput mass-spec analyses provide evidence that Y288 and Y111, both located in the dimerization regions of the protein (Sala et al., 2017a), can be phosphorylated (www.phosphosite.org), and phosphomimetic mutations of these tyrosines to glutamic acid (E) make testin mechanosensitive. Interestingly, we found that the alanine (A) mutants are also mechanosensitive, while the phenylalanine (F) mutants are not. These data suggest that the loss of the aromatic ring in the E and A mutants impacts dimerization and frees the mechanosensitive LIM domains to recognize SFSSs. This does not entirely rule out phosphorylation as a mechanism, since phosphorylation has been shown to affect the ability of tyrosines to engage in aromatic stacking interactions, thereby impacting proteins’ local structure and function (Nishi et al., 2011; Feng et al., 2010). It is thus possible that phosphorylation of Y288 or Y111 similarly alters testin’s conformation and hence mechanosensitivity. The differences in kinetics we see between mutations at positions Y288 and Y111 are likely related to their local effect on the conformation of the protein and its associated potential impact on dimerization. Structural studies, however, will be necessary to confirm and elucidate these underlying mechanisms. It is, however, remarkable that all the mechanosensitive variants of testin showed increased localization to FAs. This is consistent with our previous observation that the regions where SFs are coupled to FAs likely contain sites of strained actin filaments (Oakes et al., 2017), though it does not preclude that testin could be binding other FA proteins. A similar tension-dependent recruitment of testin to focal adherens junctions has been observed in endothelial cells, raising the possibility that testin plays an analogous role in cell-cell junctions (Oldenburg et al., 2015).

Our data also indicate that activation of RhoA causes WT testin to relocate to SFs and recognize SFSSs (Fig. 6). This suggests that signaling proteins downstream of RhoA potentially affect testin’s biochemical and/or conformational state thereby changing its mechanosensitive behavior as well. Although our data do not exclude the possibility that SF localization of testin in the presence of CA RhoA is partially mediated by its N-terminal domains, our laser ablation analysis demonstrates that the recognition of a SFSS by FL testin under these conditions is mediated by its LIM domains (Fig. 6F). Specifically, the increase in actin signal in Fig. 3D following ablation indicates the N-terminal domains are unable to recognize SFSSs and merely colocalize with condensing actin in the regions adjacent to the strain. In contrast, the loss of actin intensity following ablation in Fig. 6F is indicative of the induction of a SFSSs (Smith et al., 2013, 2011) and similar to the actin traces we observed for all the other mechanosensitive testin variants in this study. Together, this demonstrates that SFSS recognition by FL testin in the presence of activated RhoA is mediated by its LIM domains. While previous reports have established that many downstream effectors of RhoA, including kinases, show increased activity in response to mechanical forces (Lessey et al., 2012; Torsoni et al., 2005), the precise underlying mechanisms of how RhoA activation modulates testin’s mechanosensitivity remain to be explored.

Together our results imply that testin is only associated with strained SFs under certain conditions and thus we propose that its mechanosensitivity is actively regulated by the cell. Furthermore, our findings raise the possibility that similar regulatory mechanisms apply to other LDPs, of which only a subset have been identified as mechanosensitive. For example, LIMD1, in contrast to its LIM domains, is incapable of recognizing tensed SFs (Sun et al., 2020), suggesting that its mechanosensitivity is likely also context dependent. These insights may also help clarify the functional role of testin in cells. Testin is known to affect actin-driven mechanical processes, including cell spreading, cell migration and proliferation (Griffith et al., 2004; Li et al., 2016; Sala et al., 2017b). Connecting testin’s mechanosensitivity to RhoA activity provides important context for interpreting testin’s role in these and other cellular processes. Future research will be required to assess how regulation or dysregulation of testin’s mechanosensitivity affects these many cellular functions and diseases that it is implicated in. It remains clear, however, that regulation of LDPs represent an intriguing and broad mechanism for cells to control mechanotransduction activity.

## Abbreviations

CA: constitutively active
CR: cysteine rich
DN: dominant-negative
FA: focal adhesion
FHL: four- and-a-half LIM
FL: full-length
HFF: human foreskin fibroblast
LDP: LIM domain protein
LIM: Lin-11, Isl-1, Mec-3
MEF: mouse embryonic fibroblast
PET: Prickle, Espinas, Testin
SF: stress fiber
SFSS: stress fiber strain site

## Acknowledgements

We thank the Beach lab at the Loyola University Chicago, Maywood, IL for the many helpful discussions. We thank the Ampe lab at the University of Gent for their critical input and certain testin constructs. We thank Dr. Evelyne Friederich and Dr. Elisabeth Schaffner-Reckinger from the University of Luxembourg for certain testin constructs. We are also thankful to the Beckerle lab at the University of Utah, Salt Lake City, UT for providing us with the zyxin antibody, mcherry-zyxin construct and both WT and zyxin^(-/-)^ MEFs. This work was supported in part by a National Science Foundation CAREER Award (#2000554) to P.W.O. The authors declare no competing financial interests.

## Author Contributions

S.S. and P.W.O conceived the study, performed and analyzed all experiments, and wrote the manuscript.

## Materials and methods

### Mammalian expression vectors and cloning

For expression as GFP-fusion proteins, cDNAs encoding full-length (FL) testin (amino acids 1-421), full-length Y288A (Y288A substitution), NT (CR and PET domains, amino acids 1-233), LIM 1-2-3 (amino acids 231-421), LIM 1 (amino acids 234-299), LIM 2 (amino acids 299-354) and LIM 3 (amino acids 361-421) were cloned into the multiple cloning site of pEGFP-N3 or pEGFP-C3 (Clontech) as previously described (Sala et al., 2017a; b). For expression as mApple-fusion proteins, cDNA encoding full-length testin (amino acids 1-421) and NT (amino acids 1-233) were amplified from the full-length testin pEGFP-N3 vector (Forward primer: 5’-CCGCTCGAGCTATGGACCTG-3’, Reverse primer FL: 5’-CGGGATCCCTAAGACATCCTCTTC-3’, Reverse primer NT: 5’-CGGGATCCCTATTGAGTTCTTTTGTGCTC-3’) and cloned into the mApple-C1 vector (Addgene plasmid # 54631) using the XhoI and BamHI restriction sites. The Y111E substitution was introduced into the full-length testin pEGFP-N3 vector (Forward primer Y111E: 5’-TACAGTTACCGAAGAGTGGGCTC-3’, Reverse primer Y111E: 5’-TTGATGGAGACATTCTTC-3’) using the Q5 site-directed mutagenesis kit (E0554S, New England Biolabs) to generate a point mutant. Y111A, Y111F, Y288E and Y288F point mutants of GFP-fused full-length testin were a kind gift of Dr. Evelyne Friederich and Dr. Elisabeth Schaffner-Reckinger (University of Luxembourg, Luxembourg). mCherry-zyxin was a kind gift of Dr. Mary Beckerle’s lab (University of Utah, Salt Lake City, UT). pcDNA3-EGFP-RhoA-T19N (Addgene plasmid # 12967) and pcDNA3-EGFP-RhoA-Q63L (Addgene plasmid # 12968) were kind gifts from Gary Bokoch. mApple-C1 (Addgene plasmid # 54631), mApple-actin (Addgene plasmid # 54862), mApple-paxillin (Addgene plasmid # 54935) and GFP-actin (Addgene plasmid # 56421) vectors were a kind gift from Michael Davidson. All constructs were verified by sequencing.

### Cell culture and transfection

Human foreskin fibroblasts (HFF) were obtained from ATCC (CRL-2522™). Mouse embryonic fibroblasts (MEF) and zyxin^(-/-)^ MEFs were a kind gift of Mary Beckerle’s lab (University of Utah, Salt Lake City, UT). HFFs and MEFs were cultured in Dulbecco’s Modified Eagle Medium (MT10013CV, Corning) supplemented with 10% fetal bovine serum (MT35-010-CV, Corning) and 1% antibiotic-antimycotic solution (MT30004CI, Corning) at 37°C and 5% CO_2_. 24 hours prior to each experiment, HFFs were transfected with 5 μg total DNA using a Neon electroporation system (ThermoFisher Scientific) and plated on polyacrylamide gels (traction force microscopy) or glass coverslips (laser photoablation, quantification of focal adhesion localization). 48 hours prior to laser photoablation, zyxin^(-/-)^ MEFs were transfected with 5 μg total DNA using the Fugene® 6 transfection reagent (E2691, Promega) and plated on glass coverslips.

### Live cell imaging

Cells were imaged in culture media supplemented with 20 mM HEPES (SH3023701, HyClone) at 37°C, with or without 1µM SiR-Actin (CY-SC001, Cytoskeleton) or 2 nM Calyculin A (101932-71-2, Cayman), on a Marianas Imaging System (Intelligent Imaging Innovations) consisting of an Axio Observer 7 inverted microscope (Zeiss) attached to a W1 Confocal Spinning Disk (Yokogawa) with Mesa field flattening (Intelligent Imaging Innovations), Phasor photomanipulation unit (Intelligent Imaging Innovations), motorized X,Y stage (ASI) and Prime 95B sCMOS (Photometrics) camera. Illumination was provided by a TTL triggered multifiber laser launch (Intelligent Imaging Innovations) consisting of 405, 488, 561, and 637 lasers, using 63X 1.4 NA Plan-Apochromat objective (Zeiss). Temperature and humidity were maintained using a Bold Line full enclosure incubator (Oko Labs). The microscope was controlled using Slidebook 6 Software (Intelligent Imaging Innovations).

### Laser photoablation and quantitative analysis of stress fiber strain site recruitment

Prior to initiating a timelapse, a 5µm linear region was drawn in Slidebook over the SFs that were to be damaged. Cells were imaged for 3 minutes and 30 seconds alternating between the actin and testin channels with images taken approximately every 2 seconds. After 30 seconds of imaging the steady state, the marked 5µm region was illuminated with the 405 laser at a power of 370 µW for 1.5 seconds to induce a SFSS via photoablation. The remainder of the timelapse was then imaged.

Images were analyzed in python following a scheme laid out graphically in Fig. S1A. Each timelapse was first broken into two stacks representing the testin channel and the actin channel. Each stack was first flat field corrected and photobleach corrected (Payne-Tobin Jost and Waters, 2019). The testin channel was then registered using the whole image via an efficient sub-pixel registration algorithm (Guizar-sicairos et al., 2008). The calculated registration shifts from the testin channel were then applied to the actin channel, and both channels were cropped to a region of 121 x 121 pixels (∼21 µm x 21 µm) centered on the ablation region. For each channel, an average intensity image was created by averaging the frames in the stack prior to the ablation event. A relative difference was determined at each time point by subtracting the reference image from a given frame, and then dividing by the reference image. A mask of the 5% brightest points relative to the reference image in the testin channel was created for each time point. The average value of the masked points in each channel was then plotted as a function of time to create a trace of the normalized fluorescence intensity for each movie.

Average traces were created by averaging the traces from multiple movies and plotting the mean +/- the std deviation for each time point. To calculate the *t*_1/2_ value the average trace curve starting from *t* = 0 was fitted to the following equation: *I*(*t*) = *A* − *Be*^*(−t/k)*^, where 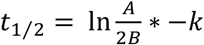. Fits are shown in Fig. S4 and calculated fit parameters for each testin construct can be found in Table S1.

The offset seen in the testin channel signal trace (e.g. Fig. 1C) is a product of making a mask of the 5% pixels that are brightest compared to the reference image. Because the location of the strain site is dependent on the local architecture of the actin cytoskeleton, we could not simply choose a predetermined region of interest to measure recruitment. By using the pixels that increased in intensity the most relative to the reference image, we were able to create dynamic masks that could identify recruitment independent of the location within the ablation region. Figs. S1B,C show how that offset changes as the number of pixels used in the mask is increased. The trace in the actin channel does not show such an offset because the masked pixels were chosen based on the testin image. For the traces in the control experiments using zyxin (Figs. 1D,H), the masked points were chosen using the zyxin channel and then applied to the testin and actin channels.

To create the spatial probability maps, we took the masks for a given movie and summed them across the entire stack (Fig. S1D). The resulting image was then rotated to align the direction of the ablation horizontally and then averaged across all the movies for a given condition. The following number of ablations were performed for each condition: FL – 16; LIM 1-2-3 – 14; LIM 1 – 15; LIM 2 – 16; LIM 3 – 14; NT – 15; Y111A – 15; Y111E – 15; Y111F – 18; Y288A – 15; Y288E – 15; Y288F – 16; FL + CA RhoA – 6; Zyxin + LIM 1-2-3 – 6; Zyxin + FL – 5; Zyxin (-/-) + LIM 1-2-3 – 6.

### Traction force microscopy and analysis

Traction force microscopy was performed as described previously (Sabass et al., 2008; Oakes et al., 2017). Coverslips were prepared by incubating with a 2% solution of 3-aminopropyltrimethyoxysilane (313255000, Acros Organics) diluted in isopropanol, followed by fixation in 1% glutaraldehyde (16360, Electron Microscopy Sciences) in ddH_2_0. Polyacrylamide gels (shear modulus: 16 kPa – final concentrations of 12% Acrylamide (1610140, Bio-Rad) and 0.15% Bis-acrylamide (1610142, Bio-Rad)) were embedded with 0.04 µm fluorescent microspheres (F8789, Invitrogen) and polymerized on activated glass coverslips for 1 hour at room temperature. After polymerization, gels were rehydrated for 45 minutes and coupled to human plasma fibronectin (FC010, Millipore) for 1 hour at room temperature using the photoactivatable cross-linker Sulfo-Sanpah (22589, Pierce Scientific). Following fibronectin cross-linking, cells were plated on the gels and allowed to spread overnight. The next day, images were taken of both the cells and underlying fluorescent beads. Number of cells imaged per condition: Fig 6B - (GFP: 35, CA GFP-RhoA: 27), Fig 6D (14). Following imaging, cells were removed from the gel using 0.05% SDS and a reference image of the fluorescent beads in the unstrained gel was taken.

Analysis of traction forces was performed using code written in Python according to previously described approaches (Sabass et al., 2008; Hanke et al., 2018). Prior to processing, images were flat field corrected and the reference bead image was aligned to the bead image with the cell attached. Displacements in the beads were calculated using an optical flow algorithm in OpenCV (Open Source Computer Vision Library, https://github/itseez/opencv) with a window size of 8 pixels. Traction stresses were calculated using the FTTC approach (Butler et al., 2002; Sabass et al., 2008) as previously described with a regularization parameter of 6.1 x 10^-4. The strain energy was calculated by summing one half the product of the strain and traction vectors in the region under the cell (Oakes et al., 2014) and normalized by the cell area as measured using the GFP image of the cell.

### Quantification of FA localization

Analysis was performed using custom code written in Python and outlined graphically in Fig. S5. Briefly, thresholding of the GFP-testin intensity images was performed to create a cell mask. mApple-paxillin intensity images were flat field-corrected and a Laplacian of Gaussian filter was applied for edge detection and creation of a FA mask. FAs were dilated using a 10×10 square structuring element and multiplied by the inverted cell mask to create a cytoplasmic donut-shaped region surrounding the FAs. The FA mask and surrounding donut mask were used to calculate the ratio (FA/cytoplasm ratio) of the average intensity in the corresponding regions of the testin intensity image. The average FA/cytoplasm ratio was then calculated per cell. Per GFP-testin variant, a total of *n* cells was analyzed (WT: 49, Y111A: 54, Y111E: 52, Y111F: 52, Y288A: 53, Y288E: 52, Y288F: 44).

### Western blotting

Cells were lysed for 20 min on ice in lysis buffer containing 50 mM Tris HCl (pH 7.6), 125 mM NaCl, 5% glycerol, 0.2% NP-40 (ab142227, Abcam), 1.5 mM MgCl_2_ and protease and phosphatase inhibitor cocktail (78441, Thermo Scientific). Cell lysates were analyzed using SDS-PAGE and 10% polyacrylamide gels, and transferred to a Whatman Protran nitrocellulose membrane (1620115, Biorad). Membranes were blocked for 1 hour at room temperature in Odyssey blocking buffer (92740000, LI-COR). After blocking, membranes were incubated at 4 °C overnight with primary antibody in Odyssey blocking buffer. Subsequently, membranes were washed 3 times in PBS containing 0.1% Tween 20 (BP337-100, Fisher Scientific) for 10 min and incubated for 1 hour at room temperature with secondary antibody in Odyssey blocking buffer. Finally, membranes were washed 3 times in PBS/Tween 20 for 10 min and rinsed in Milli-Q-H_2_O. The Odyssey® Classic infrared imaging device (LI-COR) was used for signal detection. The following primary antibodies were used: mouse anti-GAPDH (MA5-15738, Thermofisher, 1/3000 dilution) and rabbit anti-zyxin (ABD1463, Millipore, 1/1000 dilution) which was a kind gift of Mary Beckerle’s lab (University of Utah, Salt Lake City, UT). The following LI-COR secondary antibodies were used at a 1/10000 dilution: goat anti-rabbit IRdye800 (925-32211), goat anti-mouse IRdye680 (925-68070).

### Statistical analysis

Statistical analyses were performed in Python using the nonparametric multiple comparison Kruskal−Wallis one-way ANOVA test with Dunn’s posthoc method. Details about sample size and p-values are included in the Materials and Methods section and figure legends.

### Software

All images were exported from Slidebook as 16-bit TIFF files and analyzed in python. All custom code is available at (https://github.com/OakesLab).

**Table S1:** Subcellular localization and recruitment magnitude/kinetics of the testin variants used in this study.

| Testin variant | Subcellular localization | Stress fiber Strain site | Amplitude (fold increase) | Plateau max (fold increase) | T<sub>1/2</sub> (s) |
| --- | --- | --- | --- | --- | --- |
| FL | Cytoplasm/(FA) | / | n/a | n/a | n/a |
| NT | SF | / | n/a | n/a | n/a |
| LIM 1-2-3 | FA | ✓ | 2.30 +/- 0.12 | 2.26 +/- 0.06 | 10.87 +/- 1.83 |
| LIM 1 | FA | ✓ | 1.48 +/- 0.09 | 1.35 +/- 0.05 | 6.39 +/- 1.33 |
| LIM 2 | FA | ✓ | 1.55 +/- 0.12 | 1.46 +/- 0.09 | 10.55 +/- 2.31 |
| LIM 3 | FA | ✓ | 1.23 +/- 0.11 | 1.03 +/- 0.04 | 6.47 +/- 1.30 |
| Y288A | FA | ✓ | 2.20 +/- 6.62 | 2.42 +/- 6.65 | 366.76 +/- 1250 |
| Y288E* | FA | ✓ | 1.26 +/- 1.46 | 1.47 +/- 1.48 | 285.86 +/- 419 |
| Y288F | Cytoplasm/(FA) | / | n/a | n/a | n/a |
| Y111A | FA | ✓ | 1.04 +/- 0.10 | 1.34 +/- 0.10 | 16.78 +/- 4.97 |
| Y111E | FA | ✓ | 0.92 +/- 0.10 | 1.13 +/- 0.10 | 21.99 +/- 5.75 |
| Y111F | Cytoplasm/(FA) | / | n/a | n/a | n/a |

\*due to the slow kinetics of this construct the fit was based off of traces that were ~4.5 minutes in length compared to ~3 min in length for all other constructs

**Supplementary figure 1:**
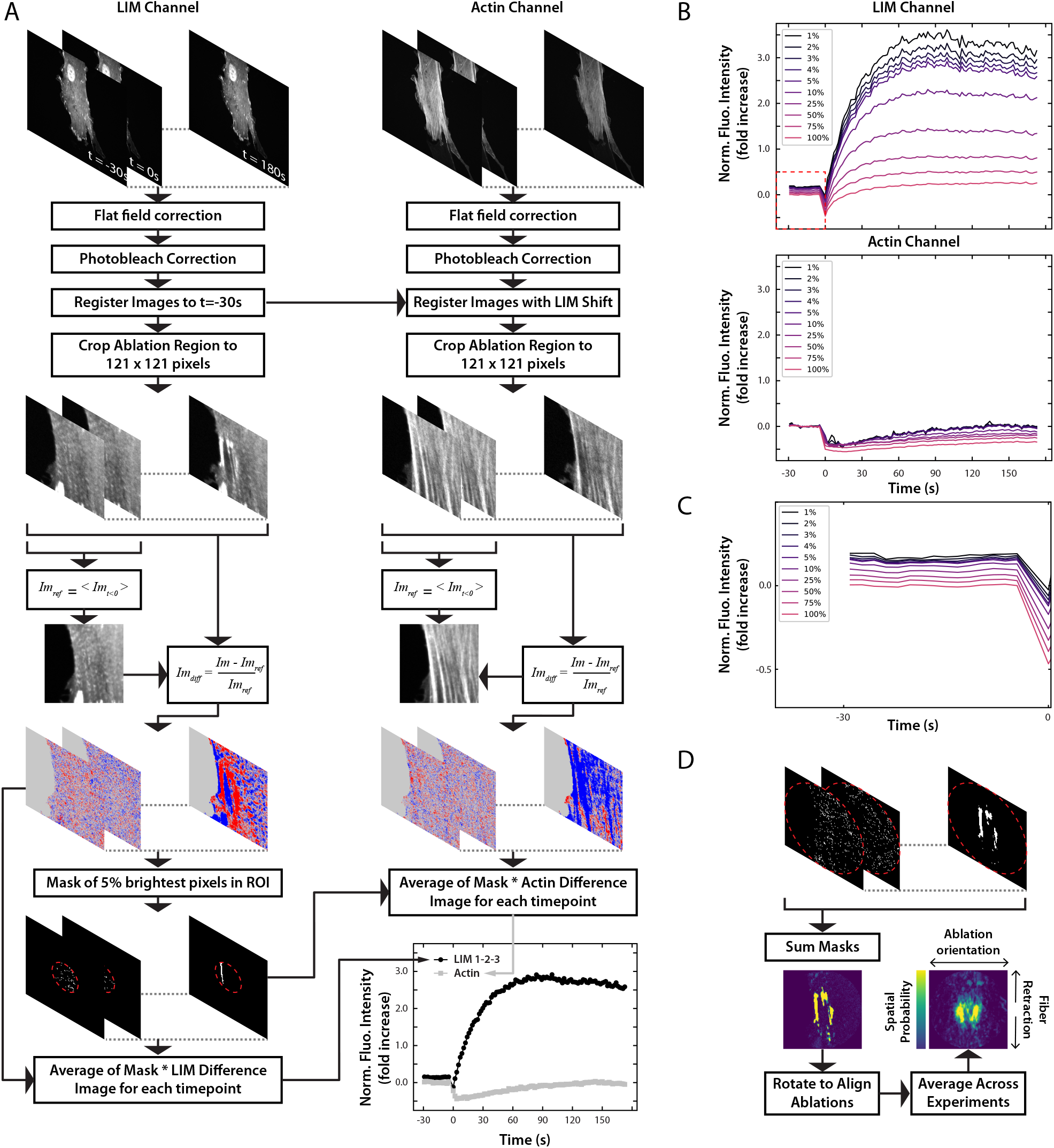
Laser photoablation analysis. **A)** A graphical representation of the workflow for analyzing the laser ablation and recruitment of testin to SFSSs. Masks were created based off the testin channel for all experiments (except Figs. 1D,H where zyxin was used instead) and used to calculate the intensity of both the testin and the actin intensities. **B)** The effect of using a different percentage of pixels to create the mask regions causes an offset from zero. **C)** The workflow to calculate the spatial probability maps.

**Supplementary figure 2:**
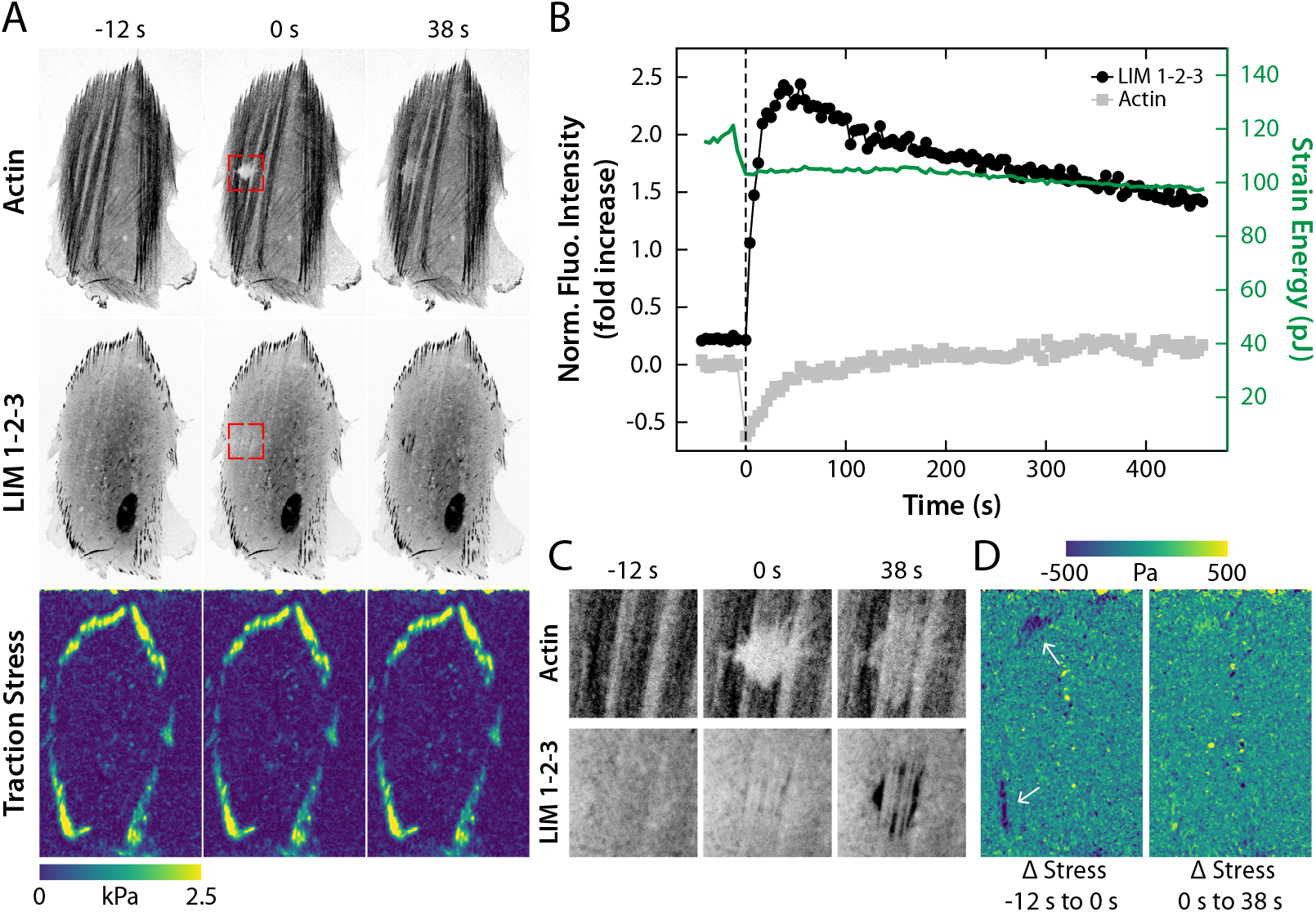
The LIM domains of testin recognize strained stress fiber. **A)** HFFs expressing mApple-actin and GFP-LIM 1-2-3 and the corresponding traction stresses. **B)** Simultaneous measurements of the total cellular strain energy and relative LIM 1-2-3 and actin intensities at the site of strain indicate the ablated SFs remain under tension. **C)** Magnified inset images (red boxes in **A**) display the localization of each construct during the timelapse (Movie S2). **D)** Difference in traction stresses pre-ablation and post-ablation show that traction stresses are reduced at the ends of the SFs (white arrows) that are damaged during ablation

**Supplementary figure 3:**
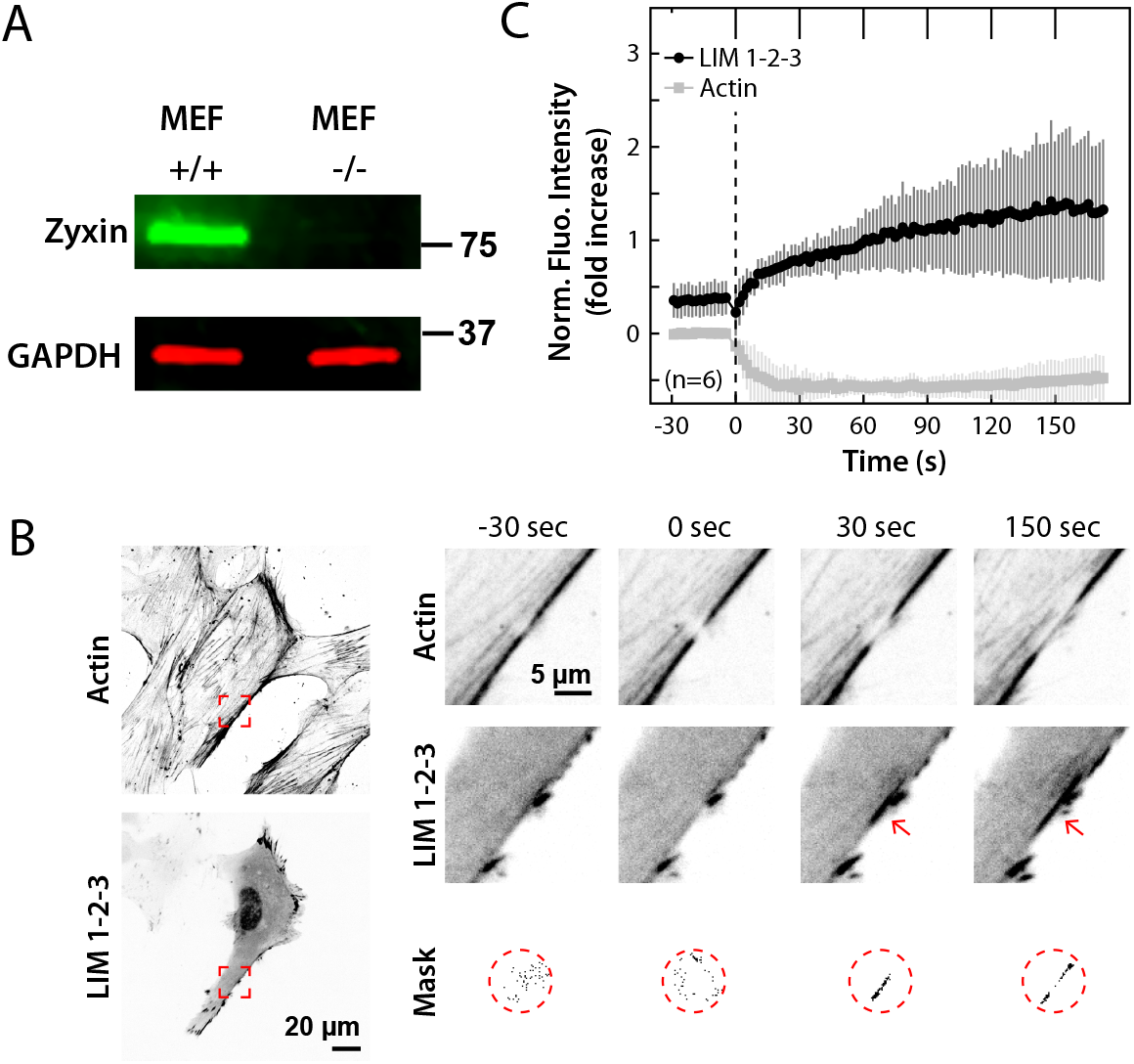
SFSS recognition by testin’s LIM domains is independent of zyxin. Western blot analysis of WT (MEF +/+) and zyxin KO (MEF -/-) MEF lysates showing no detectable zyxin levels in the zyxin^(-/-)^ MEFs. **B)** GFP-LIM 1-2-3 expressing zyxin^(-/-)^ MEFs stained with SiR-Actin. Magnified inset images (red boxed regions) display the localization of LIM 1-2-3 and actin signal during timelapse showing recruitment of LIM 1-2-3 to SFSSs (red arrows, Movie S4). Masks show the pixels in the region surrounding the ablation (red dashed circle) that increased the most in intensity compared to their pre-ablation state. **C)** Average fluorescence intensity trace and standard deviation of the LIM 1-2-3 and actin signals in the region of strain. Dashed vertical line indicates the ablation timepoint.

**Supplementary figure 4:**
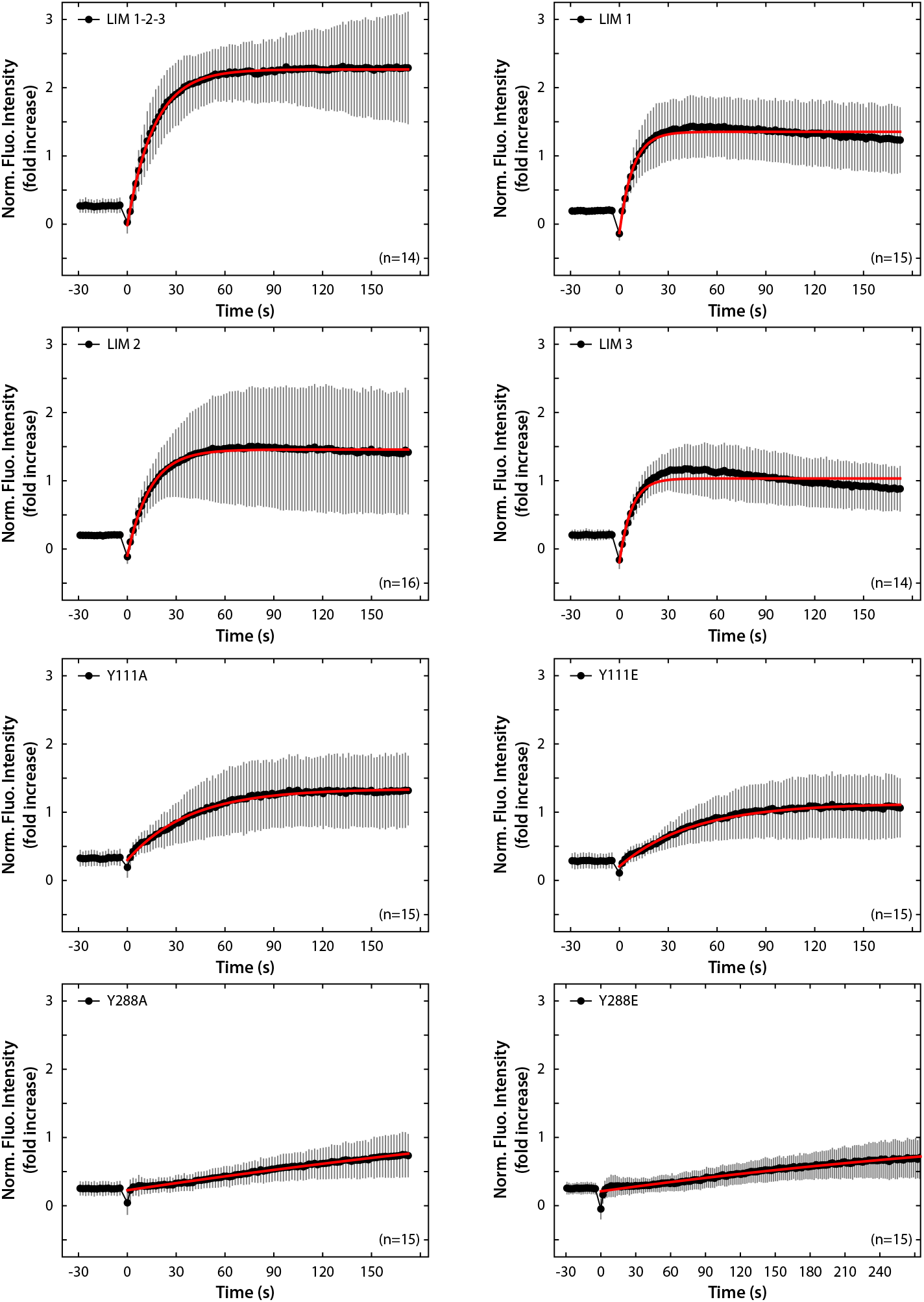
Exponential curve fitting of the average fluorescence intensity traces of mechanosensitive testin variants. Average fluorescence intensity traces are shown for the mechanosensitive testin variants found in this study. The average fluorescence intensity (thick black line) and standard deviation are plotted for each testin variant. An exponential model (red line) with the following equation was used to fit to the average traces: *I*(*t*) = *A* − *Be*^*(−t/k)*^ with A: maximum plateau value, B: amplitude, k: time constant. Note that curve fitting for the Y288E mutant is based on a 280-seconds time period for fitting purposes.

**Supplementary figure 5:**
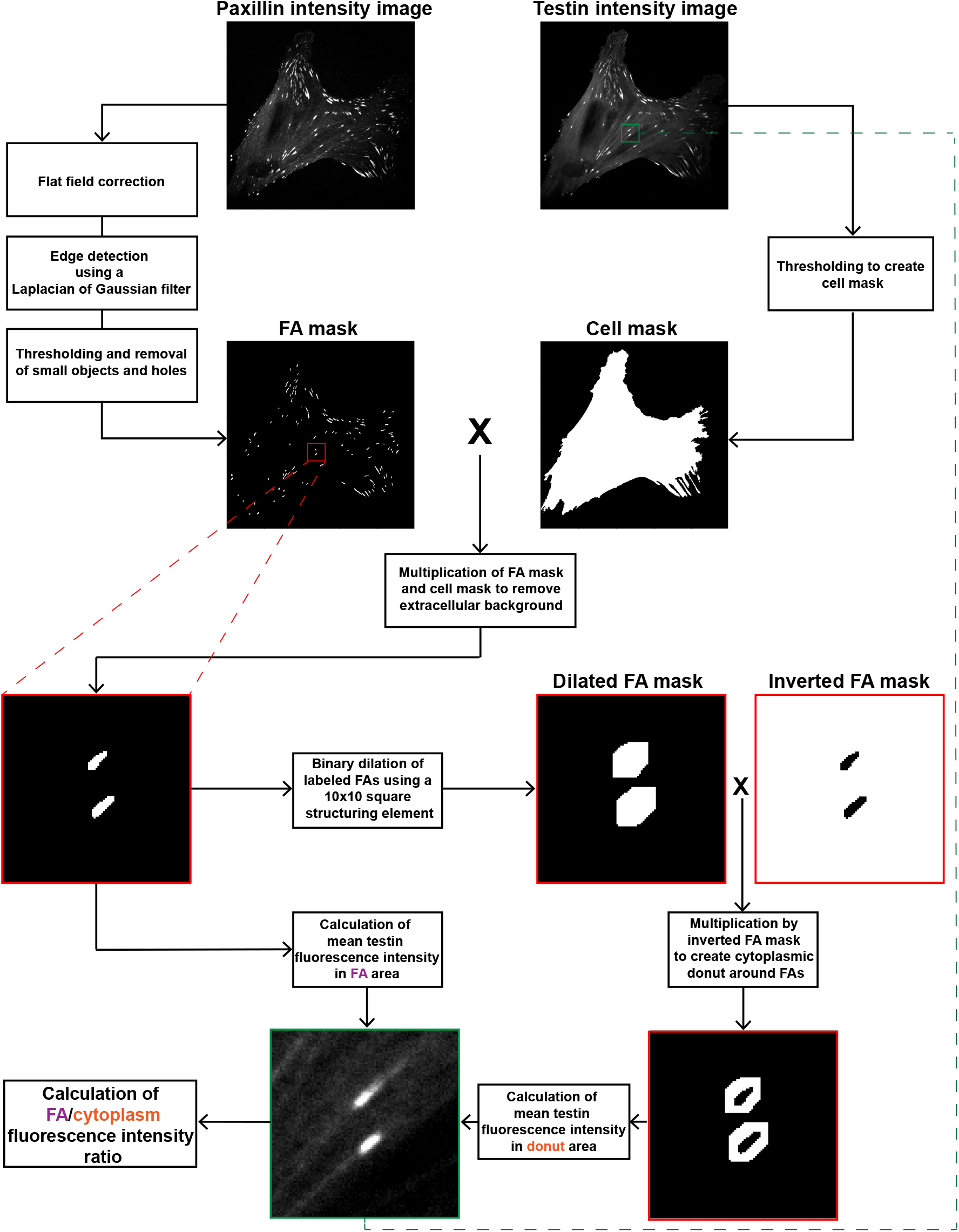
Focal adhesion localization analysis workflow. Each GFP-testin mutant (testin intensity image) investigated in Fig. 4 and Fig. 5 was co-expressed in HFFs with mApple-paxillin (paxillin intensity image) which was used as an FA marker in the analysis. Flow chart illustrates the different manipulations performed on the input intensity images resulting in the calculation of the ratio of the fluorescence intensity in the FA region and surrounding cytoplasmic region (donut) for a given GFP-testin variant.

**Movie S1:** The LIM domains of testin, but not the full-length or N-terminal half of the protein, recognize SFSSs.

**Movie S2:** Photodamaged SFs are not fully severed and remain under tension while recruiting the LIM domains of testin.

**Movie S3:** Testin’s LIM domains, but not the full-length protein, colocalize with zyxin at SFSSs.

**Movie S4:** Testin’s LIM domains relocate to SFSSs in a zyxin independent manner.

**Movie S5:** The LIM domains of testin recognize strained but not severed SFs.

**Movie S6:** Each individual LIM domain of testin relocates to SFSSs.

**Movie S7:** The N-terminal domains and C-terminal LIM domains of testin recognize different mechanical states of SFs.

**Movie S8:** The Y111A/E and Y288A/E point mutants of testin relocate to SFSSs.

**Movie S9:** RhoA drives testin to SFSSs.

